# Predicted UV-C Sensitivity of Human and Non-human Vertebrate (+) ssRNA Viruses

**DOI:** 10.1101/2021.05.10.443521

**Authors:** Brahmaiah Pendyala, Ankit Patras

**Author notes:** Correspondence: Co-Corresponding authors: Ankit Patras, Ph.D. Associate Professor; Brahmaiah Pendyala, Ph.D. Research Scientist /6018.

## Abstract

Epidemic and pandemic infectious diseases caused by RNA viruses constitute a significant hazard to human and animal health. Disinfection is an essential aspect of infection prevention and control measures. In this study, we estimated UV-C sensitivity of 83 human and veterinary pathogenic (+) ssRNA viruses by developed pyrimidine dinucleotide frequency-based genomic model. The data showed that the avian infectious bronchitis virus (genus: γ-coronavirus) with an estimated D_90_ value of 17.8 J/m^2^ was highly UV sensitive, whereas Salivirus NG-J1 (genus: salivirus) with a D_90_ value of 346.4 J/m^2^ was highly UV resistant. Overall, the trend of UV-C sensitivity of (+) ssRNA virus families followed as Coronaviridae < Flaviviridae < Togadoviridae < Arteriviridae, Matonaviridae, Astroviridae < Caciviridae < Picornaviridae < Nodaviridae < Herpeviridae. The results revealed that the enveloped viral families (Coronaviridae, Flaviviridae, Togadoviridae Arteriviridae, and Matonaviridae) are more UV-C sensitive than other nonenveloped families. Further validation of the model estimated UV sensitivity with literature available experimental data showed good agreement of predicted values. The estimates presented here could make it possible to reasonably predict UV-C disinfection efficiency of human and veterinary pathogenic viruses, which need specific biosafety requirements and/or difficult to cultivate in lab conditions.

## Introduction

Ten virus families whose members are pathogenic to humans and animals possess positive-sense (+) single-stranded (ss) RNA genomes. The families Astroviridae, Caliciviridae, Picornaviridae, Nodaviridae and Hepeviridae are characterized by non-enveloped, whereas other families Coronaviridae, Flaviviridae, Togaviridae, Arteriviridae and Matonaviridae have enveloped capsids (https://viralzone.expasy.org/294). The spread and persistence of these pathogenic viruses in diverse environments, such as hospitals, residential, public areas, pet care facilities, animal sheds, animal husbandry, etc., emphasize developing efficient decontamination processes to control epidemic and pandemic outbreaks (Cozad and Jones, 2003). Conventional chemical decontamination procedures are time-consuming, labor and resource-intensive, prone to high degrees of human error, and not applicable to air disinfection (McGinn et al., 2020). Alternative physical disinfection methods, such as germicidal ultraviolet light treatment, have gained importance due to their potential to disinfect air (Reed, 2010) and overwhelm the limitations mentioned above (McGinn et al., 2020).

The germicidal ultraviolet light disinfection method uses UV-C light to disinfect microorganisms by damaging the nucleic acids, causing them to be unable to replicate and alters vital cellular functions (Patras et al., 2020). It is well known that the disinfection level of microorganisms by UV-C light depends on their UV susceptibility, defined as D90 or D10 (dose for 90% inactivation or 10% survival) expressed as J/m2 or mJ/cm2 (Patras et al., 2020). UV-C sensitivity of a wide range of microorganisms has been reported, including vegetative and spore forms of bacteria, yeast, fungi, protozoa, algae, and viruses (Malayeri et al., 2016; Gopisetty et al., 2019; Pendyala et al., 2019, 2020a, 2021). However, the UV-C sensitivity data for many (> 80 %) human and animal pathogenic viruses is not available due to the prerequisite for biosafety level (BSL)-3 containment and the need for specifically trained skilled labor and cultivation limitations in the laboratory environment (Pendyala et al., 2020b). Acquiring the knowledge of UV susceptibility of target viruses is essential to deliver sufficient doses for efficient decontamination of the environment.

Our earlier study developed and validated a genome-sequence-based mathematical model (*r^2^ =* 0.90) to predict the UV sensitivity and identify potential SARS-CoV-2 and human norovirus surrogates (Pendyala et al., 2020b). This model was developed based on the pyrimidine dinucleotides frequency (PyNNF) of genome sequence, that effects the formation of pyrimidine dimers and 6-4 photoproducts and thereby UV susceptibility. The objective of the study was to estimate the UV-C sensitivity of 83 human and veterinary pathogenic viruses, belongs to all the families of (+) ssRNA viruses (Coronaviridae, Flaviviridae, Togadoviridae, Arteriviridae, Matonaviridae, Astroviridae, Caciviridae, Picornaviridae, Nodaviridae, Herpeviridae) by using developed pyrimidine dinucleotide frequency based mathematical model. Further validation of the model-predicted data by comparison with literature available experimental data.

## Materials and Methods

### Collection and determination of genomic parameters; genome size, and calculation of pyrimidine dinucleotide frequency value (PyNNFV)

We collected the genomic sequence of (+) ssRNA viruses belonging to families of *Flaviviridae, Picornaviridae, Arteriviridae, Coronaviridae, Togaviridae, Retroviridae, Astroviridae, Calciviridae, Nodaviridae, Hepeviridae*, and *Matonaviridae*. The size and nucleotide sequences of genomes used in this study were directly obtained from the available NCBI genome database (Table 1–5). A novel R code was developed to count the PyNNs by the exclusive method (each pyrimidine considered in one PyNN combination only) in the order of TT > TC > CT > CC and considered 100% probability when PyNN are flanked by pyrimidine on both sides and 50% probability for PyNN flanked by purine on either side. Further PyNNFV values were calculated by the following equation (Pendyala et al., 2020b).

**Table 1:** Genomic model predicted UV-C (254 nm) sensitivity (D_90_) of Flaviviridae family viruses.

| **Virus** | **Genus** | **Host** | **Disease** | **NCBI #** | **Genome (bp)** | **PyNNFV** | **$D_{90}$ (J/m<sup>2</sup>)** |
| --- | --- | --- | --- | --- | --- | --- | --- |
| Bovine viral diarrhea virus 1 | Pestivirus | Mammals | Hemorrhagic syndromes, abortion, fatal mucosal disease | NC_001461.1 | 12573 | 0.000793 | 26.3 |
| Usutu virus | Flavivirus | Human, birds, mosquitoes | Encephalitis | NC_006551.1 | 11066 | 0.001518 | 40.7 |
| Murray Valley encephalitis virus | Flavivirus | Human, mosquitoes | Encephalitis | NC_000943.1 | 11014 | 0.001139 | 33.2 |
| Japanese encephalitis virus | Flavivirus | Human, horses, birds, mosquitoes | Encephalitis | NC_001437.1 | 10976 | 0.001262 | 35.6 |
| West Nile virus | Flavivirus | Human, birds, ticks, mosquitoes | Encephalitis | NC_001563.2 | 10962 | 0.001374 | 37.9 |
| Langat virus | Flavivirus | Human, ticks | Encephalitis | NC_003690.1 | 10943 | 0.001078 | 31.9 |
| Saint Louis encephalitis virus | Flavivirus | Human, birds, mosquitoes | Encephalitis | NC_007580.2 | 10940 | 0.001004 | 30.5 |
| Louping ill virus | Flavivirus | Human, mammals, ticks | Encephalitis | NC_001809.1 | 10871 | 0.001117 | 32.7 |
| Tick-borne encephalitis virus | Flavivirus | Humans, small rodents, ticks | Encephalitis | DQ401140.3 | 11097 | 0.001141 | 33.2 |
| Yellow fever virus | Flavivirus | Human, monkeys, mosquitoes | Hemorrhagic fever | NC_002031.1 | 10862 | 0.001489 | 40.2 |
| Powassan virus | Flavivirus | Human, ticks | Encephalitis | NC_003687.1 | 10839 | 0.001205 | 34.5 |
| Zika virus | Flavivirus | Human, monkeys, mosquitoes | Fever, joint pain, rash | NC_012532.1 | 10794 | 0.001194 | 34.3 |
| Dengue virus 1 | Flavivirus | Human, mosquitoes | Hemorrhagic fever | NC_001477.1 | 10735 | 0.001015 | 30.7 |
| Dengue virus 2 | Flavivirus | Human, mosquitoes | Hemorrhagic fever | NC_001474.2 | 10723 | 0.000935 | 29.1 |
| Hepatitis C virus 2 | Hepacivirus | Human | Hepatitis | NC_009823.1 | 9711 | 0.004857 | 107.5 |
| Hepatitis C virus 1 | Hepacivirus | Human | Hepatitis | NC_004102.1 | 9646 | 0.004986 | 110.0 |
| GB virus C/Hepatitis G virus | Pegivirus | Human | None | NC_001710.1 | 9392 | 0.004022 | 90.8 |

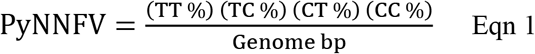

### Estimation of UV-C sensitivity (D_90_) values

Calculated PyNNF values were used to estimate the UV-C sensitivity of viruses using the reported linear regression model with r^2^ = 0.90 (Eqn 2) from our previous study (Pendyala et al., 2020b).

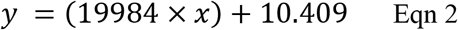

## Results and discussion

Table 1–5 depicts the selected viruses of various families of (+) ssRNA pathogenic viruses, and genus, host, diseases caused, NCBI GenBank ID, genome size, and estimated PyNNFV and D_90_ values. The genome size values ranged from 4528 bp to 30033 bp, and PyNNFV varied from 0.000371 to 0.016812. Graphical representation of data shows the estimated D_90_ values of different virus families were in the order of Coronaviridae < Flaviviridae < Togadoviridae < Arteriviridae, Matonaviridae, Astroviridae < Caciviridae < Picornaviridae < Nodaviridae < Hepeviridae (Figure 1). The results revealed that the enveloped virus families (Coronaviridae, Flaviviridae, Togadoviridae Arteriviridae and Matonaviridae) are more UV-C sensitive than other nonenveloped families.

**Figure 1:**
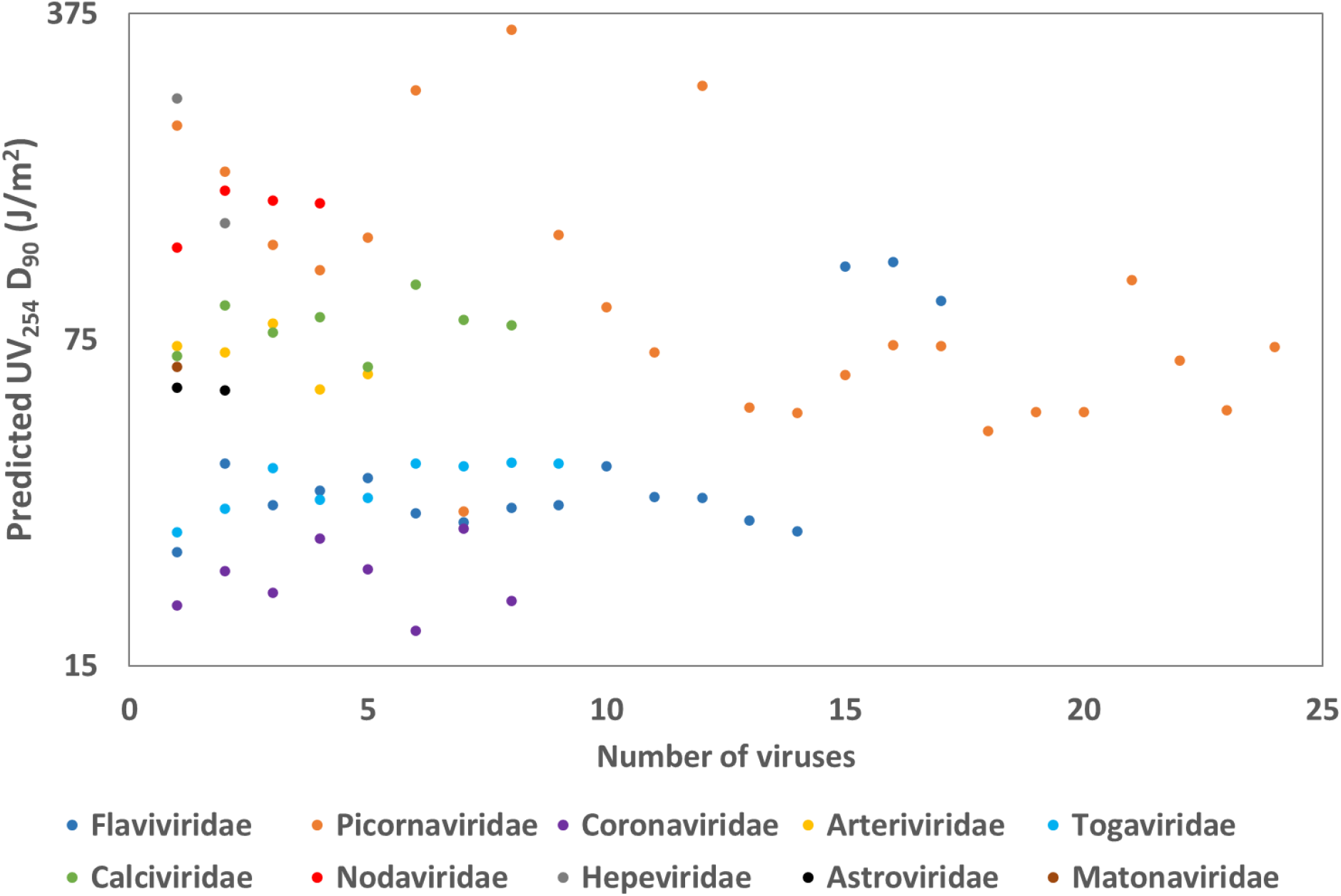
Predicted UV_254_ D_90_ values of (+) ssRNA vertebrate virus families

## Elucidation of the variability of calculated UV-C sensitivity (D_90_) at the genus level

### Flaviviridae

The estimated UV-C sensitivity of different genera of the flaviviridae family was shown in Table 1. The data shows the genus pestivirus had higher UV-C sensitivity with D_90_ of 26.3 J/m^2^than flavivirus (29.1 - 40.7 J/m^2^), pegivirus (90.8 J/m^2^), and hepacivirus (107.5 - 110 J/m^2^).

### Picornaviridae

Table 2 shows the predicted UV-C sensitivity of various genera of the picornaviviridae family. Bluegill picornavirus belongs to limnipivirus, was predicted to be highly UV sensitive with D_90_ of 33.2 J/m^2^, whereas salivirus NG-J1 was highly UV resistant (346.4 J/m^2^). The results revealed that the D_90_ values of major genus enterovirus and genera of hepatovirus, tremovirus, sapelovirus, avihepatovirus, avisvirus, and cosavirus was in the range of 47.9 – 88 J/m^2^. Medium UV sensitivity (100.9 – 125.7 J/m^2^) was observed with genera teschovirus, erbovirus, rosavirus, dicipivirus, and cardiovirus. Suboptimal UV resistance with D_90_ 171.9 to 267.7 J/m^2^was noticed with genera megrivirus, sicinivirus, kobuvirus, and sakobuvirus.

**Table 2:** Genomic model predicted UV-C (254 nm) sensitivity (D_90_) of Picornaviridae family viruses.

| Virus | Genus | Host | Disease | NCBI # | Genome (bp) | PyNNFV | $D_{90}$ (J/m<sup>2</sup>) |
| --- | --- | --- | --- | --- | --- | --- | --- |
| Siccinivirus 1 | Siccinivirus | Chicken, birds | - | NC_023861.1 | 9276 | 0.010280 | 215.8 |
| Turkey hepatitis virus | Megrivirus | Turkey | Hepatitis | NC_021201.1 | 9075 | 0.008082 | 171.9 |
| Rosavirus 2 | Rosavirus | Human | - | NC_024070.1 | 8931 | 0.005468 | 119.7 |
| Equine rhinitis B virus 1 | Erbovirus | Horse | - | NC_003983.1 | 8828 | 0.004771 | 105.7 |
| Canine picodisticistrovirus | Dicipivirus | Dog | - | NC_021178.1 | 8785 | 0.005690 | 124.1 |
| Aichi virus | Kobuvirus | Human | Gastroenteritis | NC_001918.1 | 8251 | 0.012327 | 256.7 |
| Bluegill picornavirus | Limnipivirus | Fish | - | NC_018506.1 | 8050 | 0.001090 | 32.2 |
| Salivirus NG-J1 | Salivirus | Human | Gastroenteritis | NC_012957.1 | 7982 | 0.016812 | 346.4 |
| Encephalomyocarditis virus | Cardiovirus | Human, mouse, rat, pig | Encephalitis | NC_001479.1 | 7835 | 0.005768 | 125.7 |
| Human cosavirus | Cosavirus | Human | - | NC_023984.1 | 7802 | 0.003883 | 88.0 |
| Duck hepatitis A virus | Avihepatovirus | Ducks and geese | Fatal hepatitis | NC_008250.2 | 7711 | 0.003003 | 70.4 |
| Feline sakobuvirus A | Sakobuvirus | Cat | - | NC_022802.1 | 7507 | 0.012626 | 262.7 |
| Porcine sapelovirus 1 | Sapelovirus | Pig | - | NC_003987.1 | 7491 | 0.002167 | 53.7 |
| Hepatitis A virus | Hepatovirus | Human | Hepatitis | KP879217.1 | 7476 | 0.002093 | 52.2 (51) |
| Poliovirus | Enterovirus | Human | Poliomyelitis | NC_002058.3 | 7440 | 0.002626 | 62.9 (73) |
| Coxsackievirus | Enterovirus | Human | Meningitis, myo-carditis, paralysis | KX595291.1 | 7410 | 0.003138 | 73.1 (79) |
| Turkey avisivirus | Avisivirus | Turkey | - | KC614703.1 | 7373 | 0.003120 | 72.8 |
| Human enterovirus 68 | Enterovirus | Human | Diarrhea, neurological disorder | NC_038308.1 | 7367 | 0.001877 | 47.9 |
| Human parecho virus | Enterovirus | Human | Common cold | NC_001897.1 | 7348 | 0.002096 | 52.5 (73) |
| Human rhinovirus 14 | Enterovirus | Human | Respiratory | NC_001490.1 | 7212 | 0.002110 | 52.6 |
| Porcine teschovirus 1 | Teschovirus | Pig | Teschen disease | NC_003985.1 | 7117 | 0.004529 | 100.9 |
| Human rhinovirus C | Enterovirus | Human | Respiratory | NC_009996.1 | 7099 | 0.002872 | 67.8 |
| AEV | Tremovirus | Birds | Reduced hatching. Tremors and/or ataxia. | NC_003990.1 | 7055 | 0.002126 | 52.9 |
| Swine pasivirus 1 | Pasivirus | Pig | - | NC_018226.1 | 6916 | 0.003108 | 72.5 |

Notes: $D_{90}$ in parenthesis refers literature experimental values (Pendyala et al., 2020); AEV – Avian encephalomyelitis virus)

### Arteriviridae and Coronaviridae

The families arteriviridae and coronaviridae are assigned to the order Nidovirales. The data show the coronaviridae family viruses were more UV sensitive (D_90_ 17.8 – 28.1 J/m^2^) than arteriviridae viruses (D_90_ 58.8 – 81.2 J/m^2^) (Table 3). In coronaviridae, genus γ-coronavirus was noticed to be more UV sensitive (D_90_ 17.8 J/m^2^) and β-coronavirus (MERS coronavirus) was more UV resistant (D_90_ 28.1 J/m^2^). The λ-arterivirus and δ-arterivirus were identified as more UV sensitive and UV resistant genera in arteriviridae.

**Table 3:** Genomic model predicted UV-C (254 nm) sensitivity (D_90_) of Arteriviridae and Coronaviridae family viruses.

| **Virus** | **Genus** | **Host** | **Disease** | **NCBI #** | **Genome (bp)** | **PyNNFV** | **$D_{90}$ (J/m<sup>2</sup>)** |
| --- | --- | --- | --- | --- | --- | --- | --- |
| Equine arteritis virus | α-arterivirus | Horse | vascular lesions, fever, edema, abortion | NC_002532.2 | 12704 | 0.003112 | 72.6 |
| Lelystad virus | β-arterivirus | Pig | Abortions and respiratory disease | M96262.2 | 15111 | 0.003014 | 70.6 |
| Simian hemorrhagic fever virus | δ-arterivirus | Monkey | Fever, edema, dehydration, hemorrhages, death (almost 100%) | NC_003092.2 | 15717 | 0.003541 | 81.2 |
| Forest pouched giant rat arterivirus | λ-arterivirus | Rat | - | NC_026439.1 | 14953 | 0.002423 | 58.8 |
| Wobbly possum disease virus | κ-arterivirus | Brushtail possum | - | NC_026811.2 | 12917 | 0.002640 | 63.2 |
| Human coronavirus 229E | α-coronavirus | Human | Respiratory | NC_002645.1 | 27317 | 0.000489 | 20.2 |
| SARS coronavirus 1 | β-coronavirus | Human, bats, palm civet | Respiratory | NC_004718.3 | 29751 | 0.000674 | 23.9 |
| SARS coronavirus 2 | β-coronavirus | Human, bats, pangolin? | Covid-19 | MT192772.1 | 29891 | 0.000555 | 21.5 |
| MERS coronavirus | β-coronavirus | Human, Tomb bat | Respiratory | MH734115.1 | 30033 | 0.000883 | 28.1 |
| Thrush coronavirus HKU12-600 | δ-coronavirus | grey-backed Thrush | Respiratory | NC_011549.1 | 26396 | 0.000690 | 24.2 |
| Avian infectious bronchitis virus | γ-coronavirus | Chicken and turkey | Respiratory | NC_001451.1 | 27608 | 0.000371 | 17.8 |
| White bream virus (Toro virinae (subfamily)) | Bafinivirus | Fish | Hemorrhagic liver with necrotic areas, splenomegaly, enteritis | NC_008516.1 | 26660 | 0.000954 | 29.5 |
| Breda virus (Toro-virinae (subfamily)) | Torovirus, | Human, cattle, pig, horse | Gastroenteritis | NC_007447.1 | 28475 | 0.000517 | 20.7 |

### Togadoviridae and Matonaviridae

In this family, the genus alphavirus includes mosquito-borne human and veterinary pathogenic viruses. The model predicted UV D_90_ values were between 20.9 – 40.9 J/m^2^, minimum with O’nyong-nyong virus and maximum with Venezuelan equine encephalitis virus (Table 4). Rubella virus belongs to the genus rubivirus, and family matonaviridae had predicted D_90_ value of 65.5 J/m^2^.

**Table 4:** Genomic model predicted UV-C (254 nm) sensitivity (D_90_) of Togaviridaeae and Matonaviridae family viruses.

| **Virus** | **Genus** | **Host** | **Disease** | **NCBI #** | **Genome (bp)** | **PyNNFV** | **$D_{90}$ (J/m<sup>2</sup>)** |
| --- | --- | --- | --- | --- | --- | --- | --- |
| O'nyong-nyong virus | Alphavirus | Human, mosquitoes | Fever, joint pain | NC_001512.1 | 11835 | 0.000930 | 29.0 |
| Chikungunya virus | Alphavirus | Human, monkeys, mosquitoes | Fever, joint pain | NC_004162.2 | 11826 | 0.001108 | 32.6 |
| Eastern equine encephalitis virus | Alphavirus | Human, birds, mosquitoes | Encephalitis | NC_003899.1 | 11675 | 0.001470 | 39.8 |
| Ross River virus | Alphavirus | Human, mosquitoes, marsupials, | Fever, joint pain | NC_001544.1 | 11657 | 0.001187 | 34.1 |
| Barmah Forest virus | Alphavirus | Human, marsupials, mosquitoes | Fever, joint pain | NC_001786.1 | 11488 | 0.001194 | 34.3 |
| Mayaro virus | Alphavirus | Human, mosquitoes | Fever, joint pain | NC_003417.1 | 11411 | 0.001518 | 40.8 |
| Sindbis virus | Alphavirus | Human, birds, mosquitoes | Pogosta disease, Fever, joint pain | NC_001547.1 | 11703 | 0.001486 | 40.1 (55) |
| Venezuelan equine encephalitis virus | Alphavirus | Human, rodents, mosquitoes | Fever, joint pain | NC_001449.1 | 11444 | 0.001525 | 40.9 (55) |
| Western equine encephalitis virus | Alphavirus | Human, vertebrates, mosquitoes | Fever, joint pain | NC_003908.1 | 11484 | 0.001512 | 40.7 (54) |
| Rubella virus | Rubivirus | Human | Rubella | NC_001545.2 | 9762 | 0.002757 | 65.5 |

Notes: $D_{90}$ in parenthesis refers literature experimental values (Pendyala et al., 2020)

### Calciviridae

The estimated D_90_ of calciviridae family ranged from 65.7 – 98.6 J/m^2^, and the genera lagovirus and nebovirus predicted with minimum and maximum D_90_ values (Table 5). The model predicted D_90_ values of human norovirus (HNoV) groups GI, GII and GIV were 69.1, 89.0, and 77.6 J/m^2^, respectively.

**Table 5:** Genomic model predicted UV-C (254 nm) sensitivity (D_90_) of Astroviridae, Calciviridae, Nodaviridae, and Hepeviridae family viruses.

| Virus | Genus | Host | Disease | NCBI # | Genome (bp) | PyNNFV | $D_{90}$ (J/m<sup>2</sup>) |
| --- | --- | --- | --- | --- | --- | --- | --- |
| Human astrovirus | Mamastro virus | Human, mammals | Gastroenteritis | NC_001943.1 | 6813 | 0.002446 | 59.3 |
| Avastrovirus 1 | Avastrovirus | Turkey, birds | Gastroenteritis, liver, or kidney damages | Y15936.2 | 7003 | 0.002405 | 58.5 |
| Human norovirus GI | Norovirus | Human | Gastroenteritis | NC_001959.2 | 7654 | 0.002936 | 69.1 |
| Human norovirus GII | Norovirus | Human | Gastroenteritis | KF712510.1 | 7509 | 0.003934 | 89.0 |
| Human norovirus GIV | Norovirus | Human | Gastroenteritis | JF781268.1 | 7839 | 0.003360 | 77.6 |
| Sapovirus | Sapovirus | Human | Gastroenteritis | NC_006554.1 | 7476 | 0.003672 | 83.8 |
| Rabbit hemorrhagic disease virus-FRG | Lagovirus | Lagomorphs | Necrotizing hepatitis leading to fatal hemorrhages | NC_001543.1 | 7437 | 0.002764 | 65.7 |
| Newbury agent 1 virus | Nebovirus | Bovine | Necrotizing hepatitis | NC_007916.1 | 7454 | 0.004414 | 98.6 |
| Feline calicivirus | Vesivirus | Feline | Conjunctivitis, respiratory disease | NC_001481.2 | 7683 | 0.003627 | 82.9 (60) |
| Vesicular exanthema of swine virus | Vesivirus | Swine, sea mammals | Respiratory disease | NC_002551.1 | 8284 | 0.003508 | 80.5 |
| Nodamura virus | α-nodavirus | Mammals, fishes | Paralysis and death | AF174533.1, AF174534.1 | 4540 | 0.005406 | 118.4 |
| Striped jack nervous necrosis virus | β-nodavirus | Fish | Viral encephalopathy and retinopathy | NC_003448.1, NC_003449.1 | 4528 | 0.007331 | 156.9 |
| Dragon grouper nervous necrosis virus | β-nodavirus | Fish | Viral nervous necrosis | AY721616.1 | 4536 | 0.006953 | 149.4 |
| Barfin flounder virus | β-nodavirus | Fish | Viral nervous necrosis | EU826137.1 | 4533 | 0.006839 | 147.1 |
| Hepatitis E virus | Hepevirus | Human, pig, monkey, chicken | Hepatitis | NC_001434.1 | 7176 | 0.011850 | 247.2 |
| Cutthroat trout virus | Piscihepevirus | Fish | - | NC_015521.1 | 7410 | 0.006150 | 133.3 |

Notes: $D_{90}$ in parenthesis refers literature experimental values (Pendyala et al., 2020)

### Nodaviridae

This family viruses comprised of are pathogenic to fish, comprised of two genera: α-nodavirus and β-nodavirus. The D_90_ data show the high resistance to UV (D_90_ varies from 118.4 – 156.9 J/m^2^), α-nodavirus more sensitive than β-nodavirus (Table 5).

### Astroviridae

The estimated UV sensitivity of two genera, mamastrovirus and avastrovirus were 58.5 and 59.3 J/m^2^, respectively (Table 5).

### Hepeviridae

Hepatitis E virus belongs to genus hepevirus had predicted D_90_ value of 247.2 J/m^2^, while other genus piscihepevirus (causes disease in fish) with lower D_90_ of 133.3 J/m^2^ (Table 5).

Though the developed genomic sequence-based parameter (PyNNFV) model may be sufficient to estimate UV-C susceptibility of viruses in many scenarios, the generation of more experimental data at specific regions (where the model does not have enough empirical data) is required to improve the accuracy of our predicted D_90_ values. In conclusion, our model predicted D90 values could be helpful to develop an efficient UV-C treatment process to achieve the target disinfection level of specific (+) ssRNA viruses where experimental UV-C sensitivity data is not available or feasible.

## Note

There are no conflicts to declare

